# Computational framework for exploring the interplay of diet and gut microbiota in autism

**DOI:** 10.1101/422931

**Authors:** Meghana Venkata Palukuri, Shruti Shivakumar, Swagatika Sahoo, Raghunathan Rengaswamy

## Abstract

Autism spectrum disorder (ASD)^1^ refers to the set of complex neurological disorders characterized by repetitive behaviour. The reported occurrence of abnormal gut bacteria, along with prevalence of gastrointestinal disorders in ASD indicate its strong correlation with the gut microflora. Our study aims to understand the role of diet and gut bacteria in ASD via an integrated constraint-based and PBPK model. Genome scale models of five major gut bacteria, which were reported to be associated with ASD, were integrated with the human host, i.e., the combined small intestinal enterocyte and neuronal brain model. Simultaneously, a permeability-limited two sub compartment PBPK model was developed to determine the distribution of bacteria-derived toxins in the body. The important results include, (i) inclusion of probiotics into the diet of autistic case restores gut balance, majorly seen as a result of reduced oxidative stress in the brain and the gut, (ii) microbiome and diet together mediate host metabolism in autism, majorly via the nucleotide, central carbon, amino acid, and reactive oxygen species metabolisms, and (iii) gut bacterial-specific secretions contribute to autistic metabotype. Thus, the presented integrated model is the first ever quantitative model, providing a mechanistic basis for autism pathogenesis, capturing known biomarkers, as well as, highlighting the potential of novel dietary modifications in alleviating the symptoms of autism.

## 1. Introduction

Autism spectrum disorder refers to the group of neurodevelopmental disabilities, which includes autistic disorder and pervasive developmental disorder. The typical clinical signs include social impairment, reduced cognitive capability, restricted communication skills, and repetitive behaviour (1). Although, the development of autism has traditionally been associated with genetic and environmental factors (1,2) recent reports claim this to be present in minority (3), and analysis, PBPK: physiologically based pharmacokinetic modelling, FBA: flux balance analysis, FVA: flux variability analysis and emphasize the role of inherent metabolic disturbances (4,5), including potential diagnostic biomarkers in urine (6) and plasma (7) With a current incidence of 1 in 68, autism presents comorbid patterns specifically with gastro-intestinal symptoms and mitochondrial dysfunctions (4). Additionally, there is abnormal gut microbiome composition. While the pathogenic bacteria *Clostridium, Bacteroides* and *Desulfovibrio* are present in significantly higher concentrations in the autistic gut (8-11), beneficial *Bifidobacterium* and *Lactobacillus* bacteria are present in reduced quantities (9,12,13).

These findings further link gut microbiome with autism, via the microbe-gut-brain connections, and strongly support the leaky-gut hypothesis (8) The leaky-gut or intestinal hyperpermeability, is the widening of tight junctions in the gut wall, leading to gut epithelial cells losing the ability to discern between molecules passing from the gut to the bloodstream and vice versa. Typically, increased levels of toxins like propionic acid (released by *Bacteroides*) and lipo-polysaccharide (released by *Desulfovibrio*) trigger the production of pro-inflammatory cytokines. This increases the gut permeability, thereby, allowing the toxins into the bloodstream, which can further breach the blood-brain barrier and cause autistic characteristics (14-17). Numerous studies have also identified and linked elevated levels of reactive oxygen species (ROS) to autism (4,18,19). ROS include superoxide, peroxide, etc. that are synthesized under normal physiological conditions. While superoxide is mainly synthesized during oxidative phosphorylation in mitochondria, peroxide is encountered in amino acid metabolism (via monoamine oxidase, E.C. 1.4.3.4.), peroxisomal fatty acid oxidation (acyl CoA oxidase, E.C. 1.3.3.6.), etc. (20). Thus, it is necessary to mathematically model the complex interactions between gut microbiome and observable autistic symptoms, not only to better understand ASD pathogenesis, but also to direct future research and suggest measures to stall autism regression.

Constraint-based reconstruction and analysis (COBRA) (21) remains one of the preferred systems biology methods to study biological cell systems with mechanistic details. Numerous studies have been published that use the COBRA approach to study the relationship between gut bacteria and the host gut, giving rise to potential applications in understanding various human diseases, including metabolic disorders and cancer (22-24). In line with this, a collection of 773 metabolic reconstructions/models of 13 most abundantly found human gut microbe phyla was made available (25). However, COBRA operates on steady-state conditions (details in experimental procedure), hence, to capture the dynamics of autism pathogenesis a whole body physiologically based pharmacokinetic (PBPK) modelling (26) approach is required, specifically to model the transport and effect of gut-derived toxins.

In this paper, we present, the first ever, integrated model comprising constraint-based genome-scale metabolic networks of human gut, brain and a semi-detailed PBPK model of six organs (Figures 1 and 2). The genome-scale metabolic models of the human small intestine and the five important bacteria species (i.e., *Bacteroides vulgatus, Clostridium perfringens, Desulfovibrio desulfuricans, Lactobacillus acidophilus* and *Bifidobacterium longum*) were combined to represent the human gut. Further, the gut model was connected to the genome scale metabolic model of human brain, via the PBPK transport model. Given the model set up (Figure 2), following questions were answered: (i) can modelling host-microbe interactions under different dietary conditions hint towards dietary treatment options for autism? (ii) what are the fundamental biochemical factors driving autism?, and (iii) what is the role of bacteria-derived toxins/metabolites in autism?

**Figure 1:**
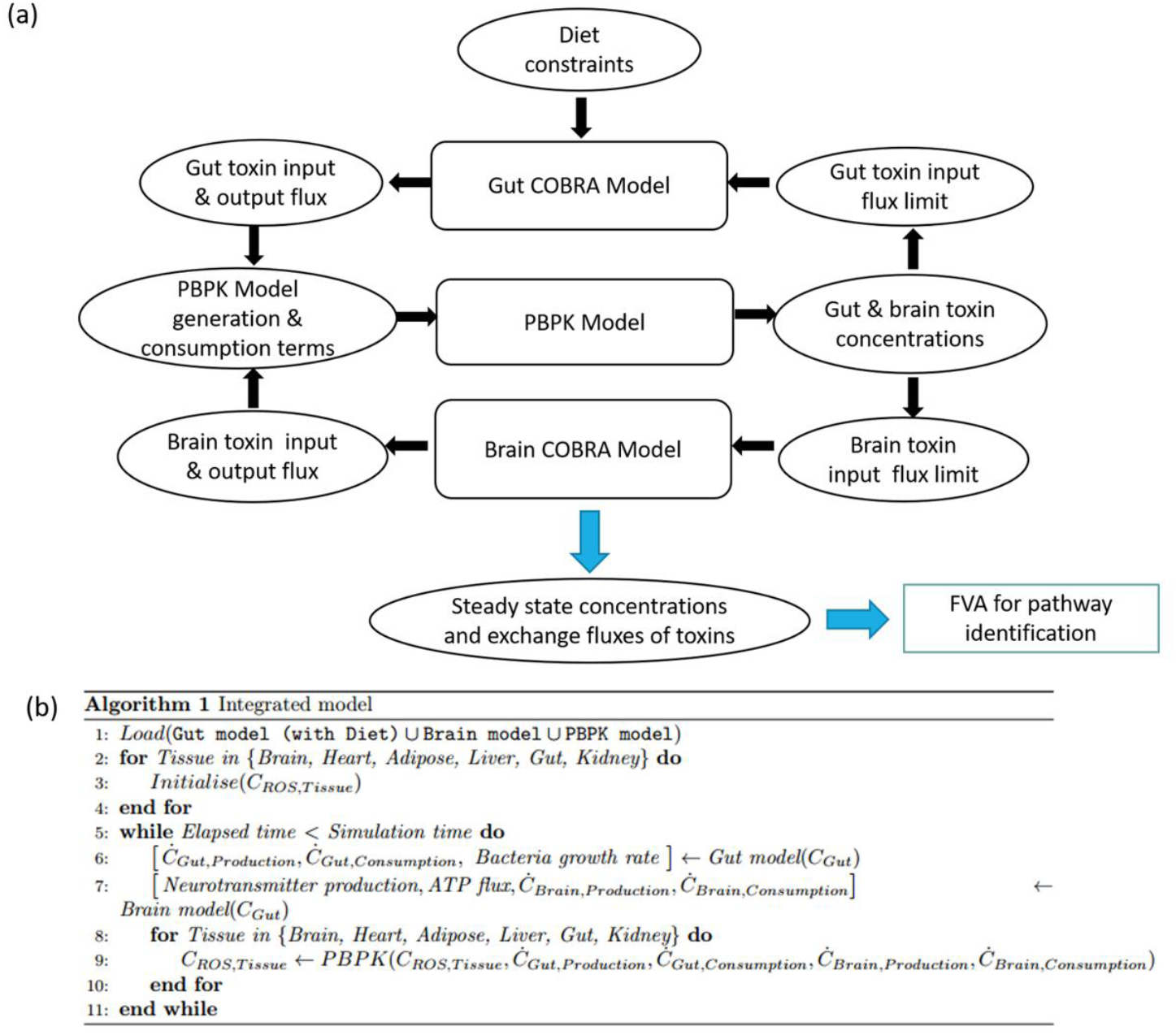
(a) The integrated model, showing the interactions between the gut, brain COBRA models and the transport PBPK model. Simulation of the integrated model yields the steady state exchange fluxes of toxins used for further analysis through FVA to identify pathways. (b) Algorithm for the integrated model.

**Figure 2:**
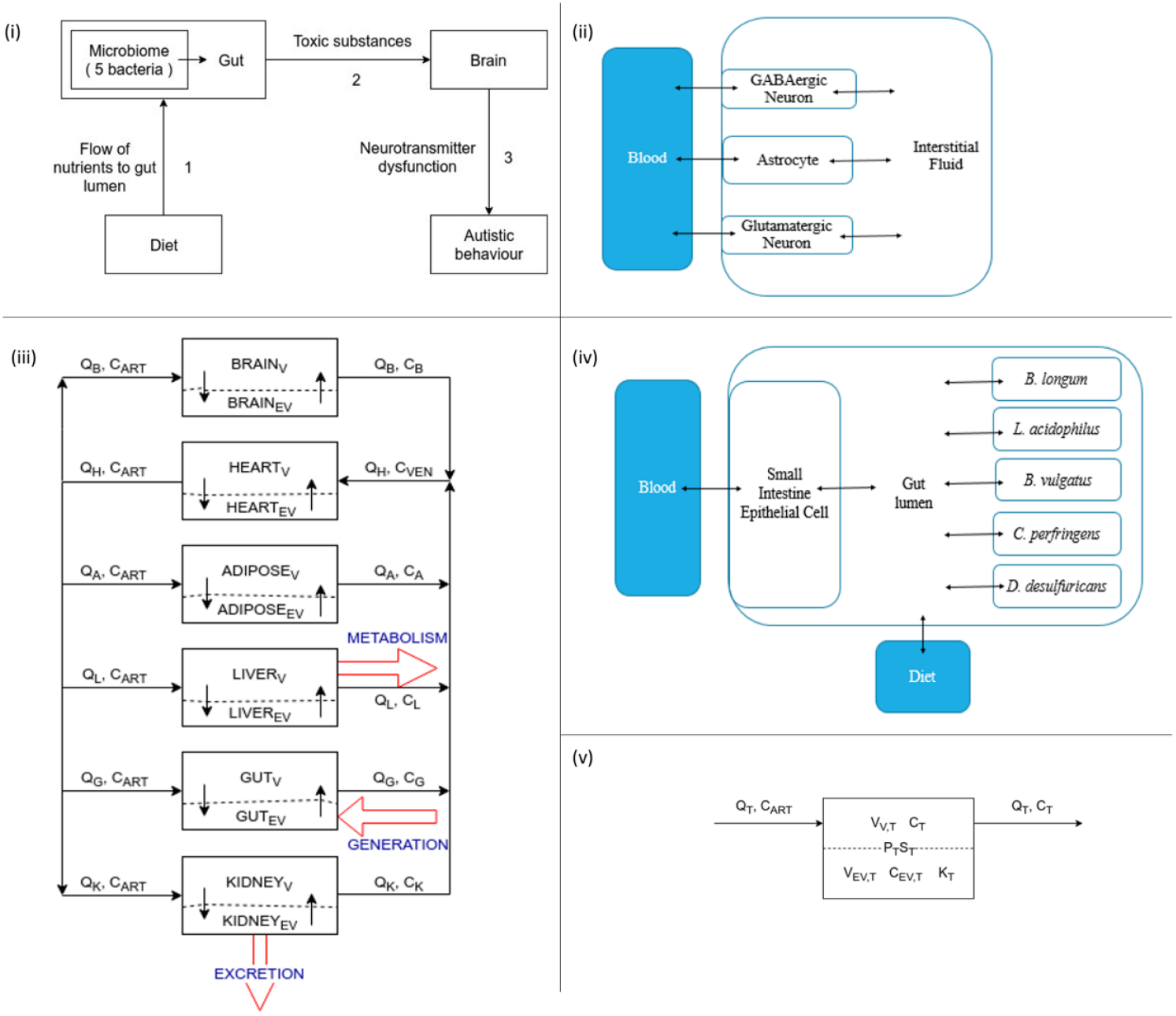
Overview of models used - (i) overall integrated model (top left) (ii) brain model and its components (top right) (iii) Whole-body PBPK model (bottom left) (iv)gut model and its components (middle right) (v) Mass balance in tissue type T with volume *V*_*T*_ modelled as permeability-limited, two sub-compartment model - a vascular sub compartment, *V* of volume *V*_*V.T*_ and an extra-vascular sub-compartment, *EV*, of volume 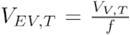 where ƒ is the tissue fractional blood volume. The molecule of interest enters the vascular region of the tissue through the arterial blood stream *ART* of flowrate *Q* and permeates across to the extra-vascular region over surface area *S*_*T*_ with permeability *P*_*T*_. It distributes according to the tissue:plasma partition coefficient, *K*_*T*_, which is the equilibrium distribution of the molecule between the two phases. The ratio between the volumes of the vascular to the extra-vascular sub-compartment is known as the tissue fractional blood volume, *f*.

## 2. Results

### 2.1. Model building

The genome-scale metabolic models of the five bacteria (25) were coupled with the genome scale model of the human small intestinal enterocyte model (27), representing the human gut, which included 7137 reactions, 6056 metabolites and 4116 genes. Further, published neuronal models (28) were combined to represent the whole-brain model, comprising of 1542 reactions, 1428 metabolites and 806 genes (Table 1). Details of the model components are given in Table S1.

**Table 1:** Individual and combined model Specifications

| Model | Number of reactions | Number of metabolites |
| --- | --- | --- |
| Bacteroides-vulgatus-ATCC-8482 | 1302 | 1110 |
| Desulfovibrio-desulfuricans-subsp-desulfuricans-DSM-642 | 1279 | 1148 |
| Clostridium-perfringens-ATCC-13124 | 1307 | 1115 |
| Lactobacillus-acidophilus-ATCC-4796 | 756 | 728 |
| Ta Bifidobacterium-longum-longum-JDM301 | 1054 | 959 |
| SIEC | 1282 | 844 |
| GABAergic neuronal model | 1068 | 993 |
| Glutamatergic neuronal model | 1067 | 993 |
| Gut microbiome – Beneficial | 3177 | 2611 |
| Gut microbiome – Harmful | 5323 | 4365 |
| Gut microbiome | 7137 | 6056 |
| Brain | 1542 | 1428 |

We developed five separate models of the gut microbiome to analyse the effect of varying gut bacteria composition, representative of their different levels in autism, i.e., (i) A ‘purely autistic’ gut (also, Gut Microbiome-Harmful) constructed by combining sIEC and the individual models of the harmful bacteria growing at maximum rate (ii) A ‘typical autistic’ gut constructed by combining the sIEC model with the individual models of all five bacteria, with the harmful bacteria forming 80% of the total gut bacterial population (iii) A ‘corrected’ gut (also, Gut Microbiome) constructed by combining the sIEC model with the individual models of all five bacteria growing at maximum rate, representing a scenario where symptoms of autism are alleviated by administering probiotics (iv) A ‘typical healthy’ gut constructed by combining sIEC and models of all five bacteria, with the beneficial bacteria forming 80% of the total gut bacterial population (v) A ‘purely healthy’ gut (also, Gut Microbiome-Beneficial) constructed by combining the sIEC model with the models of beneficial bacteria. Components of the gut models are illustrated in Figure 2, and the model details are specified in Table S1.

The PBPK model was carefully designed so as to investigate the effect of gut-derived toxins: (i) hydrogen peroxide, and (ii) superoxide on autistic symptoms, through their transport between the brain and the gut. The novelty of the PBPK model lies in its semi-detailed and highly curated nature, wherein, the physiological and biochemical parameterisation of the model was largely literature-driven. Additionally, the physicochemical parameters such as tissue:blood partition coefficient and renal clearance for each of the six organs of interest were estimated using Quantitative-Structure Property Relationships (QSPR) (29), as shown in Tables S2 and S3.

In order to understand the complex interactions between the gut microbiome composition and human host under different dietary constraints, for autistic v/s healthy case, an integrated model was constructed. The human host was represented by the COBRA genome scale metabolic models of the gut and the brain. The PBPK model was an essential connective model that was used to transport gut-derived metabolites across the COBRA genome-scale metabolic models of the tissues. Further, the PBPK model aided in setting accurate bounds for the toxin exchange reactions for the constraint based models. The importance of the integrated approach is highlighted by the fact that, in the absence of a PBPK model, the constraint based models would have been initialised with arbitrary bounds, thereby not representing true physiological conditions. We demonstrated this by choosing the steady state flux values to be 100 times the actual values obtained from the PBPK model so that we could simultaneously study the effect of increase in toxin fluxes (Table S4).

The integration of COBRA and PBPK modelling linking the two tissue types, i.e., gut and brain was carried out as detailed in Algorithm 1 (Figure 1b).

### 2.2. Model analysis techniques

Having constructed an integrated model, we now present a framework for its analysis. A multicellular model such as the gut-microbiome or the brain requires multi-objective optimization techniques for determining the growth-rates or ATP maintenance fluxes of the individual cells. Further, there were other multi-objective optimization problems posed, such as the production and consumption of various toxins and neurotransmitters, demanding equal importance to be given to each compound. Such multi-objective problems have been classically solved using pareto-optimization techniques (30,31). With this motivation, an equally weighted pareto-optimization framework using a linear search technique was developed (Figure 3). A model with varying percentages of different types of cells can be simulated by posing a multi-objective optimization problem, wherein, the growth of a cell type present in higher proportion is awarded higher weightage. Such a situation was encountered while simulating the ‘typical autistic’ and ‘typical healthy’ models, for which we developed a weighted pareto optimization technique, this time employing a binary search technique (Figure 3). The various cases for which pareto optimization were implemented are detailed in Table S5. The developed models were tested to analyse the effect of harmful and beneficial bacteria, diet, and gut-derived toxins. Two metrics were defined based on the minimum and maximum possible fluxes of reactions obtained via flux variability analysis. Highly affected pathways were then identified (Table S6). The details of the model building and analysis methods are described in experimental procedures and supporting information.

**Figure 3:**
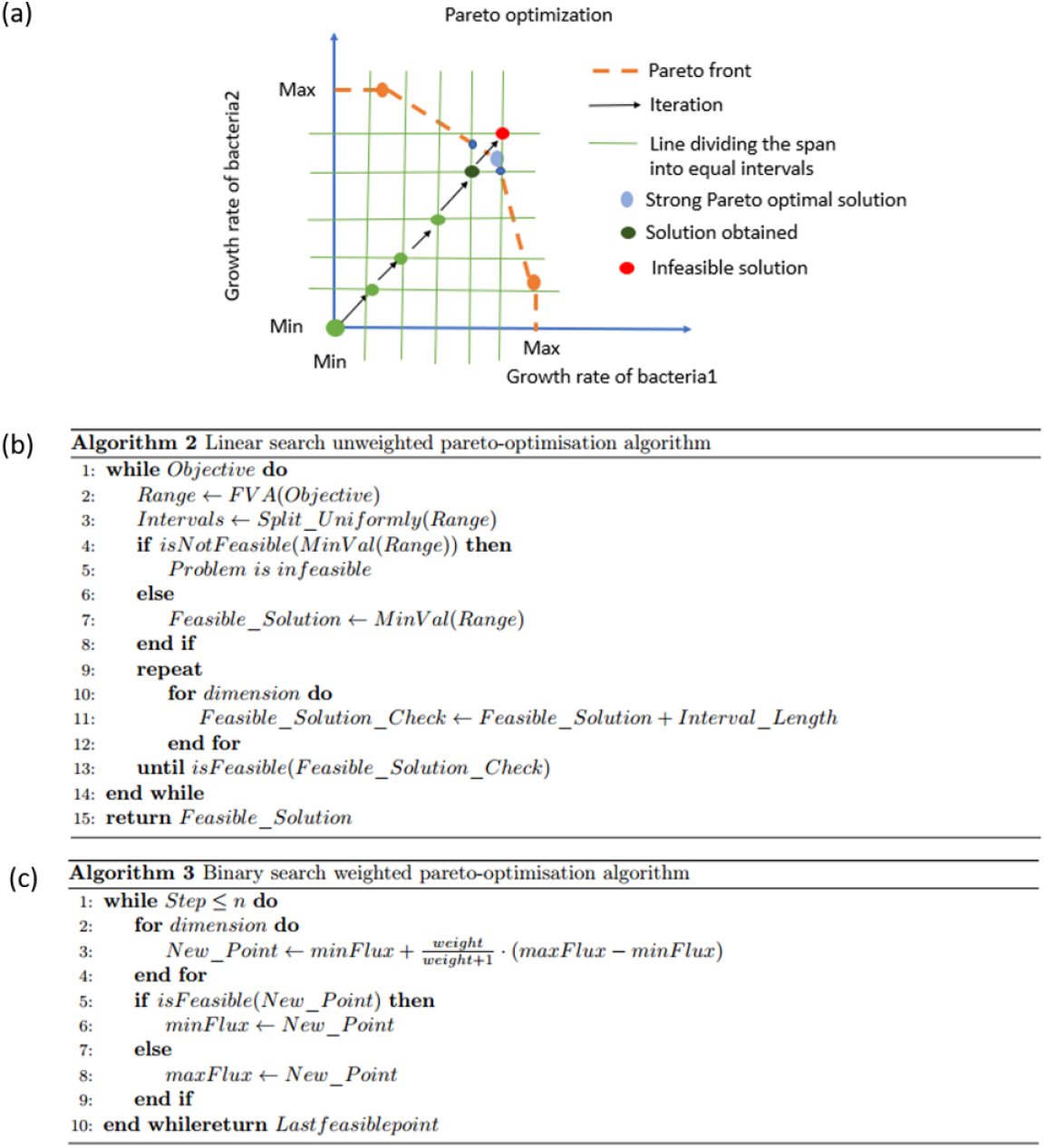
(a) Pareto optimization (linear search) (b) Linear Search weighted Pareto optimization algorithm (c) Binary Search weighted pareto-optimization algorithm.

### 2.3. Addition of probiotics improved autistic features

As mentioned earlier, the first major question of the present study was to identify the potential dietary treatment strategy for autism. Two typical dietary patterns, favouring microbial growth were chosen, i.e., a western diet and high-fibre diet (25). Each of the autistic and healthy models showed reasonable growth of each cell type under both dietary conditions (Table 2). Interestingly, we observed that the addition of the beneficial bacteria *Lactobacillus acidophilus* and *Bifidobacterium longum* to the purely autistic gut decreased the growth rates of each of the harmful bacteria, but increased the growth rates of the beneficial bacteria themselves, despite being in lesser quantity than the harmful bacteria (Table 2). This suggests that *Lactobacillus acidophilus* and *Bifidobacterium longum* can be used as probiotics to significantly lower the growth rates of harmful bacteria. However, the effect on sIEC must also be kept in mind since high dosages may be harmful, considering that sIEC growth rate decreases with increasing percentages of bacteria in general (Tables 2 and S7). This occurred due to higher competition for nutritional resources.

**Table 2:** Model functioning: (a) Growth rates of gut microbiome cells (mmol/gDW/hr)

| Model | Diet | BV | DD | CP | LA | BL | SIEC |
| --- | --- | --- | --- | --- | --- | --- | --- |
| Gut microbiome | Western | 1.05109 | 1.04102 | 0.3129 | 0.7336 | 0.9927 | 0.00972 |
|  | High Fibre | 1.06472 | 1.0585 | 0.3057 | 0.73197 | 1.0106 | 0.00933 |
| Gut microbiome - Harmful | Western | 1.2675 | 1.2220 | 0.4448 | - | - | 0.01601 |
|  | High Fibre | 1.3402 | 1.2940 | 0.4529 | - | - | 0.01601 |
| Gut microbiome - Beneficial | Western | - | - | - | 0.1586 | 0.1511 | 0.01984 |
|  | High Fibre | - | - | - | 0.3098 | 0.6589 | 0.02546 |

(b) ATP production fluxes in brain cells on consumption of Western diet (mmol/gDW/hr)
| Brain cell | ATP production |
| --- | --- |
| Astrocyte | 3.3851 |
| Glutamate neuron | 4.1379 |
| GABA neuron | 4.1398 |

An altered neurotransmission in GABAergic neurons, including loss of GABA interneurons has been identified in a subset of autism cases (1,9,32). Hence, we next simulated the effect of probiotics on brain metabolism. Maintenance flux for each of the neuronal cell types in the brain model was obtained (Table 2). Additionally, the model indicated that the flux through the synthesis reactions of GABA, an inhibitory neurotransmitter, increased on intake of beneficial bacteria (Table S8). Hence, we propose that probiotics may improve autism, majorly via increasing the availability of GABA in brain.

### 2.4. Microbiome and diet together alter intestinal and brain metabolism in autism

Metabolic aberrations in autism is a predominant feature, typically the class involving high grade gastrointestinal symptoms (4). Studies have majorly shown the association of abnormal amino acid, fatty acid, energy, and reactive species metabolism (4,5). Herein, we report that purine catabolism and nucleotide interconversion pathways are also among the most affected pathways in autism pathogenesis, when simulated under western diet (Table 3). Consequently, elevated purine catabolism, via xanthine oxidase (GeneID: 7498, *XDH,* E.C. 1.17.3.2.) was reported in autism patients (33). Further, increased uric acid levels in autistic children, and ‘anti-purinergic’ therapy has been shown to reduce autistic features in mice (34).

**Table 3:** Summary of effect of gut bacteria, diet, and reactive species metabolites on the metabolism of gut and brain. Only top ten pathways are shown. (a) Metabolic pathways affected in the gut by addition of beneficial bacteria and changing the diet from western to high-fibre. (b) Metabolic pathways affected in the gut and brain due to increased oxidative stress.

| S.No. | Pathways affected by addition of beneficial bacteria to 'purely' autistic gut | Pathways affected by change in diet |
| --- | --- | --- |
| 1 | Purine catabolism | Purine catabolism |
| 2 | Nucleotide interconversion | Glycolysis/gluconeogenesis |
| 3 | Glycolysis/gluconeogenesis | Fructose and mannose metabolism |
| 4 | Citric acid cycle | Citric acid cycle |
| 5 | Valine, leucine, and isoleucine metabolism | Pyruvate metabolism |
| 6 | Propanoate metabolism | Nucleotide interconversion |
| 7 | Pyruvate metabolism | NAD metabolism |
| 8 | Vitamin B6 metabolism | Purine synthesis |
| 9 | NAD metabolism | Vitamin B6 metabolism |
| 10 | Purine synthesis | Pyrimidine catabolism |

(b)
| S.No. | Pathways affected in the gut | Pathways affected in the brain |
| --- | --- | --- |
| 1 | Purine catabolism | ROS Detoxification |
| 2 | Citric acid cycle | Oxidative Phosphorylation |
| 3 | Alanine and aspartate metabolism | Pyruvate Metabolism |
| 4 | Purine synthesis | Glutamate metabolism |
| 5 | Vitamin B6 metabolism | NAD Metabolism |
| 6 | NAD metabolism | Citric Acid Cycle |
| 7 | Pyrimidine catabolism | Urea cycle/amino group metabolism |
| 8 | Glycolysis/gluconeogenesis | Alanine and Aspartate Metabolism |
| 9 | Arginine and Proline Metabolism | Valine, Leucine, and Isoleucine Metabolism |
| 10 | Nucleotide interconversion | Folate Metabolism |

To assess the effect of beneficial bacteria, we added these to the purely autistic gut model, and observed a higher flux through the glycolysis pathway. Contrastingly, addition of harmful bacteria, lead to higher flux in the reactions linked to oxidative stress metabolism (details in Table S9). We propose that this could be another cellular protective mechanism to combat oxidative stress, a phenomenon also seen in cancer cells (35).

Oxidative stress has been widely implicated in autism (36). In order to determine the effect of oxidative stress on metabolic pathways affected in autism, we developed and compared the ‘toxic’ model and the ‘non-toxic’ model for both the gut and the brain. The former allows high exchanges of toxins related to oxidative stress across membranes, due to the increased permeability observed in autistic conditions (37,38) while the latter corresponds to low toxin exchange. The pathways affected by oxidative stress are given in Table S4. The top affected pathways carrying more flux through its reactions in the toxic case was found to be ROS detoxification, oxidative phosphorylation and glutamate metabolic pathways.

We specifically modelled two representative ROS species, *H*_*2*_*O*_*2*_ and SOX in order to determine the effect of oxidative stress on autism. We observed that oxidative stress was reduced under the administration of probiotics, specifically when subjected to a high-fibre diet. Further, steady-state SOX levels in the brain and gut were reduced on increasing the percentage of beneficial bacteria, seen for both the diets. Lastly, steady-state SOX concentrations were found to be lower for the high-fibre diet, when compared with the western diet (Figure 4, Tables S8 - S10). Interestingly, pathways that were found to be the most affected by gut bacterial imbalance seen in autism, overlap with the pathways affected by oxidative stress (Table 3). This indicates that there is a strong metabolic inter-relationship between gut dysbiosis and cellular stress levels, majorly mediated via the amino acid, nucleotide, central carbon, and vitamin metabolisms.

**Figure 4:**
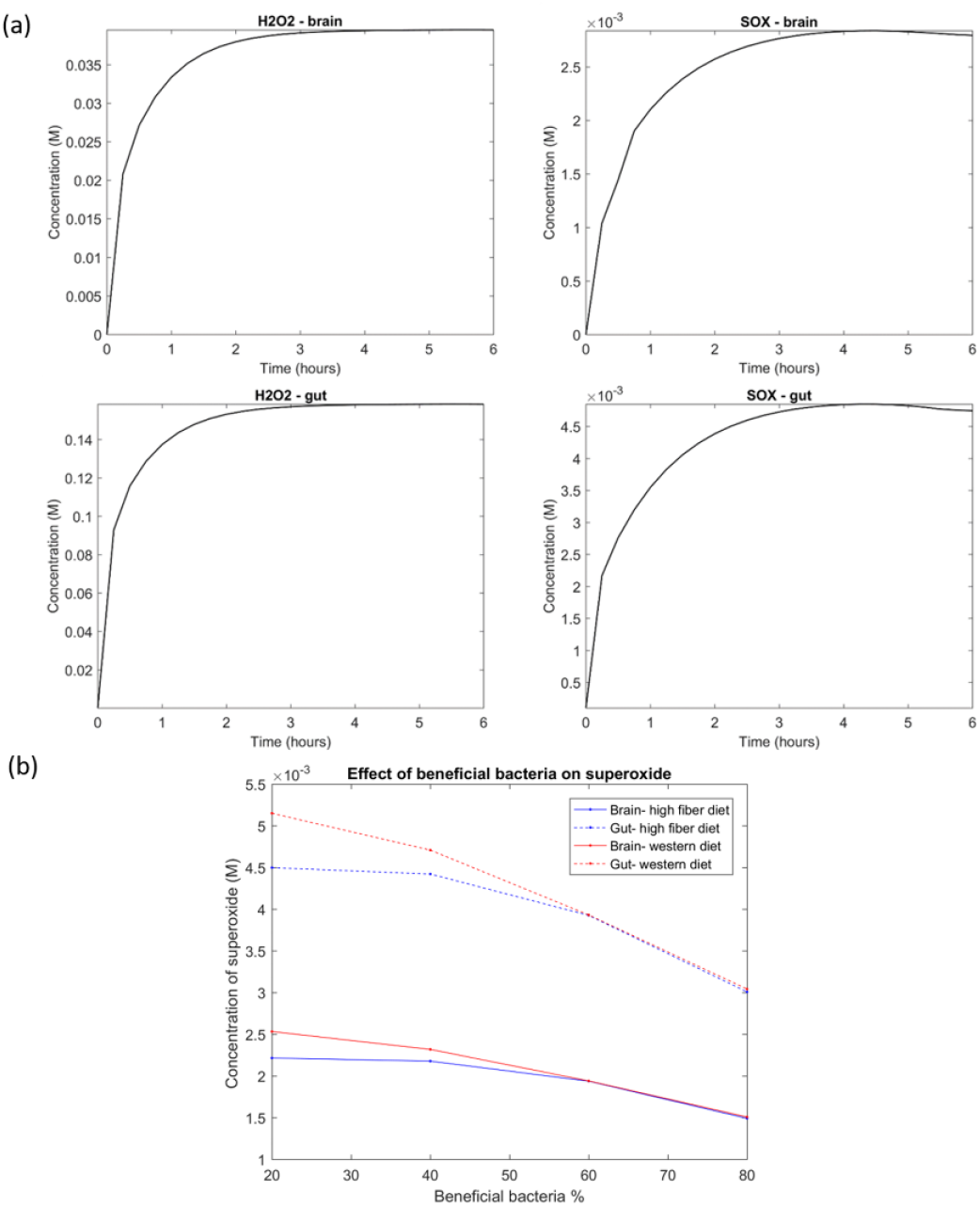
(a) Concentration Profiles of *H*_2_*O*_2_ and SOX for Gut microbiome under western diet (b) Effect of beneficial bacteria on superoxide.

We found that the concentrations of the toxins increased steadily in the organs for a small time period, after which they saturated (Figure 4). This happened, since, the initial conditions chosen were those for an ideal (or desired) case of very low toxin concentrations in the body, which would then increase naturally with the activation of metabolic pathways, producing reactive oxygen species in the gut and the brain. After a certain time period, we observed that the concentration profiles saturated, indicating that the net intake and production balanced the net consumption and excretion of ROS from the body. Further, steady state ROS concentrations (both H2O2 and SOX) were higher in the presence of harmful bacteria, indicating that the presence of harmful gut bacteria in the small intestine leads to increased production of ROS, via harmful bacteria-derived toxin interaction (Figure 4 and Table S11).

### 2.5. Bacterial secretion products

The importance of autistic-specific metabotype, i.e., secretion profile of autistic patient group was realised via recent metabolomics studies (18,39) and other screening tests (3). We predicted a total of 230 possible bacterial secretion products (Table S12). The top 15 secretion products, i.e., the exchange reactions with the highest maximum fluxes in FVA are given in Table. 4, ranked from high to low (details in Table S12). The biomarker predictive capacity of the integrated model is moderate, as it capture some of the important bacterial secretions implicated in autism, like propionic acid (15,16), lactate and pyruvate contributing to conditions linked to autism, such as mitochondrial dysfunction (40) and oxidative stress (41), as well as, ammonia (8).

We then analysed the effect of the addition of probiotics on these potential biomarkers. The flux through harmful bacteria secretions reduced as the fraction of beneficial bacteria was increased, as shown in Tables 4 and S12. Typical example include, reduced flux through ammonia excretion. Interestingly, elevated levels of ammonia have been reported to be present in autistic children (8,42).

**Table 4:**
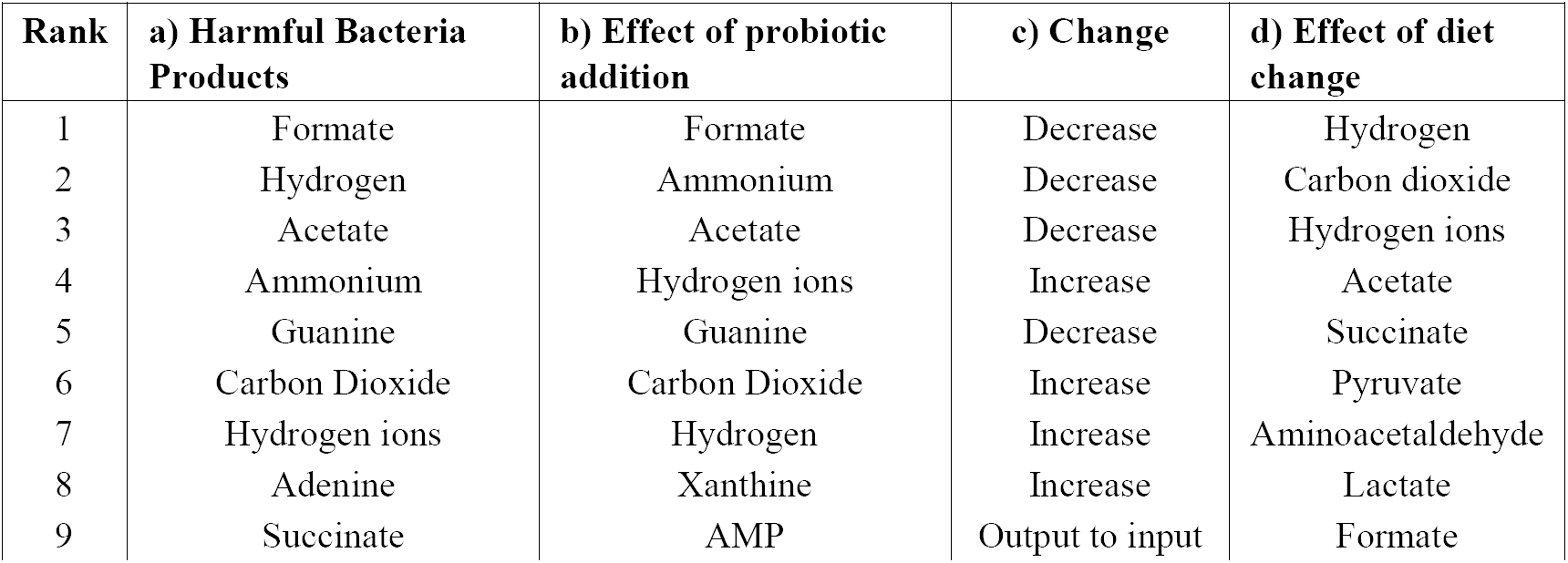
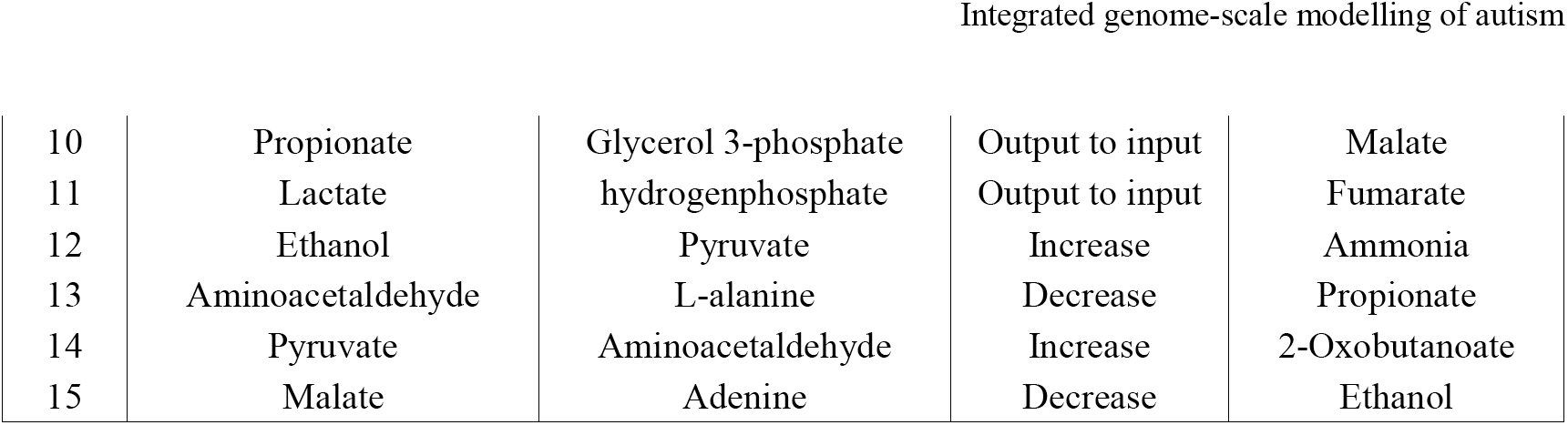
Summary of the effect of beneficial bacteria and diet on the secretion products of the harmful bacteria. Only top 15 bacterial secretion products are shown. (a) Harmful bacterial secretion products (b) Harmful bacterial secretion products that are affected by addition of beneficial bacteria, c) The change observed upon addition of the corresponding beneficial bacteria, (d) Increased secretion products after changing the diet from western to high-fibre.

Further, given these potential biomarkers, we analysed their modulation via different diets. Under western diet, the beneficial bacteria secreted 20 metabolites, 65% of which were foreign. Under high-fibre diet, the beneficial bacteria secretions increased to 25 metabolites, 68% of which were foreign. This is indicative of probiotics being relatively more effective in a high-fibre diet, when compared to a western diet.

## 3. Discussion

The presented model is the first ever representation of a hybrid modelling technique for autism, which integrates steady-state and dynamic modelling principles. The key challenge to integrating constraint-based models and PBPK models is that the former calculates fluxes (rates) of reactions, whereas the latter, computes concentrations of the metabolites of interest. This was resolved by applying the static optimization approach of dynamic FBA (43), wherein, it was assumed that the entire concentration of a metabolite was consumed by a reaction in a given time step, and thus, this maximal rate served as bounds for the input reaction.

Simulating multi-cellular systems poses several challenges. Single objective (say, biomass of a cell) optimized using FBA, maximized individual cell growth ignoring the growth of other cells, making it unsuitable for optimizing multi-cellular systems. Setting the objective function as the sum of biomass fluxes of each of the cells and optimizing using FBA yielded a solution with zero biomass fluxes for some cells and high biomass fluxes for others. Hence, pareto optimization was implemented, so as to obtain a solution with non-zero biomass fluxes for all cells (wherever possible) and to enable collective growth of the system in terms of growth of every cell.

Significant insights were obtained with respect to the effect of diet and gut bacteria in autism. Firstly, the growth of all bacteria was observed to be generally higher under the high-fibre diet when compared to western diet (Figure S1, Table 2), since fibre in the body is digested primarily by gut bacteria (44). Since the growth of healthy bacteria increased (when compared to western diet) to a larger extent than that of harmful bacteria in high-fibre diet, especially at higher percentages of healthy bacteria in the gut, we can infer that high-fibre diet is better than western diet for beneficial bacteria growth, and consequently in alleviating autistic symptoms as discussed below. Secondly, on increasing the fraction of beneficial bacteria in the gut, which is equivalent to administering probiotics to the autistic individual, we observed a monotonic decrease in the growth rate of all harmful bacteria under both the diets and a monotonic reduction in the levels of oxidative stress in both the brain and the gut (Figure 4). This integral connection between diet and gut microbes was found to majorly influence host metabolism, typically the amino acid and oxidative stress metabolism (Table 3). Consequently, our model also showed higher flux through reactions of glutamate metabolism in brain. Excessive amounts of glutamate, and an aberrant mitochondrial glutamate transporter have been implicated in autism (8,45). Additionally, the model was able to predict synergistic effects of diet and probiotics to combat autistic symptoms. These effects were further found to be modulated via gut and brain metabolic pathways. Typically the nucleotide and central carbon metabolisms in the gut and mitochondrial energy and amino acid metabolisms in the brain (Table 3).

The high-fibre diet resulted in higher flux through bacterial growth, and succeedingly in higher production of bacterial secretion products when compared to the western diet (Table 4). The model effectively captured some of the known biomarkers of autism, e.g., pyruvate and lactate. Interestingly, when analysed for bacteria-specific contribution to these secretions, *Bacteroides vulgatus* model contributed the maximum amount to the total formate and acetate generated in the autistic gut (Table 4). This makes sense, since *Bacteroides vulgatus* decomposes complex sugars and produces short-chain fatty acids (46). On the other hand, *Clostridium perfringens* model generated the maximum amount of H2. Thus, we propose that *Bacteroides vulgatus* and *Clostridium perfringens* can be the most severe drivers of autism among the implicated harmful bacteria. Further, their secretion products can be counter-balanced, leading to their significant reduction levels via addition of beneficial bacteria (Table 4). This in turn can form the basis for development of novel drugs and dietary formulations focusing on reducing the quantities of these two bacteria in order to show marked improvement in autistic symptoms.

## 4. Conclusion

We have presented a novel model building strategy, i.e., the integration of COBRA and PBPK models to identify and explain disease driving factors in autism. To this end, the integrated model of five gut microbes, a small intestinal epithelial cell connected to the brain via a transport model, was able to predict the known metabolic associations connecting the gut microbe-small intestine-brain, with considerable accuracy. Recent reports (47) have linked other bacteria to autism, i.e., *Akkermansia muciniphila* and *Sutterella wadsworthensis* that are not considered here. However, no known mechanism has been put forward for the presence of *Sutterella* in an autistic gut, as opposed to the case of *Clostridium (8).* Our model incorporates known bacteria with strong biochemical evidence for their association with autism. As strong literature evidence becomes available, scaling up with addition of new bacterial species is straightforward with the current model setup. In the current study, we have modelled increased oxidative stress by using only two representative species, i.e., SOX and H2O2. This can be extended to include other neurotoxins, through the prediction framework laid down for the PBPK model parameters. Further, our model wrongly predicted the elevation of alanine metabolism in gut and brain, as opposed to low alanine levels detected in urine samples of ASD children (18). Consequently, with inclusion of other known autism biomarkers, e.g., pro-inflammatory cytokines and bacteria-derived phenols, a wider picture of autism pathogenesis can be obtained. Nevertheless, our model can be used as a starting point for understanding both qualitative and quantitative aspects of pathogenic factors and treatment thereof, for a number of diseases that show strong association with gut microbiota, including metabolic diseases and cancer (48). We believe inclusion of diverse bacterial species, as well as, other potential organs, i.e., skeletal muscle and immune system (4) can widen the scope of mathematical modelling for autism research.

## 5. Experimental procedures

### 5.1. Model Construction

In order to explore the role of dietary and cellular metabolic factors in autism, three separate models - a combined small intestine - bacteria model representing the human gut, a combined neuronal model representing the human brain, and a PBPK transport model - were constructed and coupled together.

### 5.2. Genome scale metabolic models – gut microbiome and brain

A gut metabolic model was constructed by combining individual constraint-based genome scale metabolic models of sIEC (27), and five of the gut bacteria implicated in autism (25). These include the three ‘harmful’ bacteria, *Bacteroides vulgatus, Desulfovibrio desulfuricans* and *Clostridium perfringens;* and the two ‘protective’ bacteria - *Lactobacillus acidophilus* and *Bifidobacterium longum.* The effect of diet was analysed by introducing constraints on the model exchanges in the combined model for each of the dietary constituents. Two commonly used diets, the western diet and the high-fibre diet (25), with minor modifications (details in Table S13) were simulated.

The unconstrained gut bacteria models were obtained from the AGORA Reconstructions (25). As per intestinal conditions, exchange reactions were constrained for each of the five bacterial models. A typical example includes the aerobic condition of the small intestine, set by constraining the lower bound of oxygen exchange reactions of the bacterial models, as lb = -1mmol/gDW/hr. The constraints are detailed in Table S13.

The sIEC model, i.e., hs_sIEC611 (27) was further expanded with new pathways to capture the metabolism of bacteria-derived toxin metabolites. A total of 12 reactions and 7 metabolites were added to include metabolism of propionic acid and important reactive oxygen species i.e. *H2O*_*2*_ and SOX (details in Table S1). The gut bacteria and the sIEC interact through exchange of metabolites via the lumen compartment. Thus, to generate the combined model, 665 luminal transport reactions were added for the extracellular metabolites of the bacterium models (details in Table S14). Additionally, exchange reactions were added to the lumen compartment to allow dietary intake and excretion from the system. In order to analyse the effect of harmful bacteria on the gut, and consequently, the extent of the protective influence of the beneficial bacteria, the fractional amount of beneficial bacteria in the gut was varied from 0 to 0.8, with no beneficial bacteria representing the extreme case of a ‘purely’ autistic gut. The individual and combined model specifications are given in Table S1.

Recent investigations have shown that the neurotransmitters GABA and glutamate are implicated in autism (32,49). Thus, a whole brain model was constructed by combining the previously published models of the GABA and glutamate neurons (28). Each neuronal model comprised of two cell types - the neuron, as well as the supporting astrocyte cell. After quality checks and corrections (supporting information document), the individual neuronal models were then merged to give the combined whole-brain model.

### 5.3. PBPK transport model

The PBPK transport model considers each tissue type of interest as a two-compartment, permeability limited, well-stirred tank, consisting of a vascular compartment and an extravascular compartment. The different compartments interact with each other through the circulatory system of the body. Thus, movement and accumulation of the compound as a function of time was quantified in each tissue type through a set of ordinary differential equations. The steps involved in constructing a PBPK model are given in Table S15.

#### PBPK transport model representation

One of the main objectives of this study is to examine the effect of ROS in the brain of autistic individuals, so we included tissues that metabolize ROS, specifically SOX and H2O2, or whose function is affected by ROS in the fullbody PBPK model. Thus, the minimalistic PBPK model included (i) the heart as a source and sink of the circulatory system, (ii) the gut, (iii) the brain, as they were the central organs in our study. Since increased ROS levels are implicated in adipocyte dysfunction (50), (iv) adipose tissue was included in the model. Being the central metabolic organ, (v) the liver was included to maintain the generality of the model, and (vi) the kidneys were included to allow for excretion. In order to formulate the model equations, tissue-specific characteristics that affect the uptake of the molecule of interest, needed to be defined.

As mentioned earlier, elevated levels of ROS in the autistic gut lead to inflammation of the gut epithelial wall, which further results in increased permeability of the gut wall. It was also deduced that, once these toxins breach the blood-brain barrier, it is possible that they can cause brain inflammation (14). Thus, in order to account for the increased epithelial permeability of the gut wall and the brain in autistic individuals v/s controls, a tissue model that accounts for organ permeability in its formulation was used. Thus, a permeability-limited, two sub-compartment model was formulated and used for further analysis (Figure 2). The governing mass balance equations for this model are:

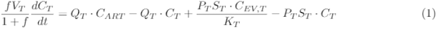

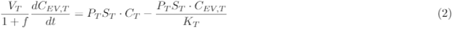

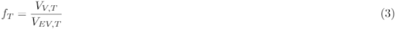

The final set of ordinary differential equations used was simply the above two mass balance equations written for each of the six tissue types, i.e., brain, heart, adipose, liver, gut and kidney (Table S16) with additional generation and consumption terms obtained from the COBRA models, as detailed in the supporting information document.

#### Parameter Estimation in PBPK model

There are three unique characteristics of a PBPK model, namely physiological, physicochemical and biochemical characteristics (Table. S17). While physiological parameters like blood flow rate through different tissue types, tissue surface area and tissue volume, and biochemical parameters like enzymatic reaction rate and Michaelis constant were largely obtained from literature, physicochemical parameters - partition coefficient *K*_*T*_, renal clearance *CL*_*R*_ and tissue permeability *P*_*T*_ - were estimated using QSPR (Tables S18 and S19).

### 5.4. The integrated COBRA-PBPK model

In the integrated model, the concentrations change at every time step, not only due to consumption or production of metabolites by the COBRA model (modelled as input and output reactions, respectively), but also, due to the rate of metabolite transport. The transport depends on permeability and other parameters that are taken into consideration by the PBPK model. This was achieved by computing the bounds on the input fluxes to the COBRA model using the concentrations obtained from the PBPK model at every time instant. To ensure input of the metabolite into the COBRA model, the reversible exchange reaction was replaced by two irreversible reactions, i.e., an input reaction constrained by the bound obtained from the PBPK model and an output reaction. The PBPK model in turn used the actual input and output exchange fluxes of the metabolite. This was computed by the COBRA analysis as its consumption and generation terms respectively, and the PBPK model computes the concentrations for the next time instant. The process repeats with the obtained concentrations, with the entire procedure described in Algorithm 1 (Figure 1b).

The COBRA simulation at each time step for the gut was constructed as follows. Under the maximal bounds set for the toxin inputs, pareto-optimal growth of all cell types of the model was computed and set as bounds for the respective biomass reactions. Now, maximal toxin production (output exchange) fluxes were determined using pareto-optimization and set as bounds to the output exchange constraints. Under these constraints, the maximal toxin input fluxes were determined, using pareto-optimisation, as the consumption terms for the PBPK model and the output exchange fluxes as the generation terms. The exact same procedure was used in the COBRA simulation for the brain at each time step, except that the ATP maintenance reaction was maximised instead of growth. Additionally, the neurotransmitter production fluxes (modelled as demand reactions for GABA and glutamate) were computed using pareto-optimization under the calculated constraints. The steps involved in the integrated model analysis is illustrated in Figure 1.

To ensure gradual intake of a metabolite by the organ, the bound on the input reaction for the next time step was set to be below 20 % of the bound at the present time step. The concentration-flux conversion is given in Equation 4.

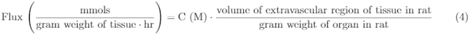

where, the volume of the extra-vascular region of the brain and the gut was obtained from the PBPK model and the gram weight were obtained from experimental values defined for rats, i.e., the small intestine was 7.26 g (51) and the brain was 2 g (52).

### 5.5. Model Analysis

#### Pareto optimization

For multi-objective maximization, a pareto optimal solution (30,31) is required. Two pareto optimization algorithms were developed and implemented, employing linear search (Algorithm 2, Figure 3) for equally weighted and binary search (Algorithm 3, Figure 3) for unequally weighted pareto-optimisation.

In a system with equal number of cells of each type, equal weighting was awarded to each objective (e.g., biomass of each cell), since it is expected that each cell in the system tries to maximize its own growth irrespective of the external conditions.

A graphical representation of the working of Algorithm 1 in the search space for the solution is given in Figure 3. Stopping criteria that can be imposed include a maximum number of iterations or the Euclidean norm of the difference between consequent iterates falling below a supplied tolerance value.

In case of weighted pareto-optimisation, a binary search technique (Algorithm 2, Figure 3) was used to obtain other pareto optimal solutions by choosing different slopes of lines from the starting point (minimum fluxes in each dimensions). For instance, consider the case where the percentage of bacterium 1 is 80%, while the percentage of bacterium 2 is 20% in the system. The algorithm will award weights in the ratio of 1:4 to the increment step in the algorithm.

#### Model comparison with FVA

FVA, which computes network variability, was used to identify the affected pathways under varying simulation conditions/ constraints. To enable comparison of the reactions of a model in terms of the effect of the change in conditions, we defined two metrics using the fluxes of a reaction in a model and its perturbed form, as given in Equations 5 and 6.

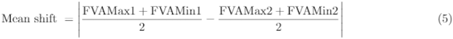

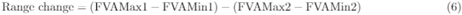

where, *FVAMax1* and *FVAMin1* were the maximum and minimum fluxes of a reaction in a model before modification of conditions. *FV AMax*2 and *FVAMin*2 are the maximum and minimum fluxes after modification of conditions.

Both these metrics were computed for each of the reactions of interest in the model and the reactions were ordered in decreasing order of mean shift. In cases of a tie, the decreasing order of range change was used to determine the order. Thus, an order was determined for the effect of modifications in a model on all the reactions of interest.

#### Pathways affected by oxidative stress

The effect of oxidative stress was determined by comparing the ‘toxic’ and the ‘non-toxic’ models, as mentioned in Section 2, for both the brain and the gut using FVA. The ‘toxic’ model allowed higher exchanges of toxins. This was obtained by setting the lower bounds of the toxin input and output exchanges to 50% of the steady state flux values obtained from the integrated model. The ‘non-toxic model’ allowed lower exchanges of toxins. This was obtained by setting the upper bounds of the toxin input and output exchanges to 50% of the steady state flux values obtained from the integrated model. The implications of the leaky-gut hypothesis and increased permeability of the blood brain barrier were examined through simulation of this phenomenon on the brain model by allowing the flow of toxins into the brain model. Further, the effect of toxins on the brain was studied by comparing a ‘toxic’ and ‘non-toxic’ model of the brain. The steady state values for the brain toxin exchanges, obtained from the integrated model, were set as the bounds on the toxin exchanges for the brain model. The same was done for the gut model. In both these analyses, the models were compared using FVA.

#### Gut - microbiome interactions and bacterial secretion products

Performing flux variability analysis, FVA (53) on the gut microbiome model and identifying the active reactions revealed that a total of 230 metabolites can be secreted and exchanged between the bacteria and sIEC models (details in Table S12). Of these, four metabolites were found to be secreted only by beneficial bacteria and 99 metabolites only by harmful bacteria. Further, 127 metabolites were secreted by both harmful and beneficial bacteria.

Thereafter, beneficial bacteria models were added to the Gut microbiome harmful models, in order to analyse the effect of beneficial bacteria on the production of secretion products from harmful bacteria. This was performed using model comparison with FVA, under western diet conditions. Additionally, the effect of diet was determined by comparing with the gut microbiome model under high-fibre diet conditions (details in Table S12).

## Acknowledgements

This work was supported by the INSPIRE faculty award, Department of Science & Technology, India (DST/INSPIRE/04/2015/000036) to SS. SS and RR also thank IIT Madras for funding for the dual degree program that supported MPV. The authors thank IBSE team for valuable discussions.

## Author contributions

MPV and Shivakumar, S. performed the experiments and analysed the results. MPV developed algorithms and analysis tools. SS and RR designed the study and planned the experiments. All the authors wrote the manuscript.

## Supporting information

Here in we provide description of the supplementary tables.

Table S1: Excel file containing details of all the developed models, i.e., gut microbiome reactions, gut microbiome metabolites, brain reactions, brain metabolites, gut microbiome-beneficial reactions, gut microbiome-beneficial metabolites, gut microbiome-harmful metabolites, and gut microbiome-harmful reactions.

Table S2: Table containing partition coefficient predicted for propionic acid, hydrogen peroxide and SOX using QSPR

Table S3: Table containing renal clearance (L/s) predicted for propionic acid, hydrogen peroxide and sulphur dioxide using QSPR

Table S4: Excel file containing details of the effect of oxidative stress, i.e., (a) comparison of brain metabolism of non-toxic model, and toxic model under steady state toxin fluxes, (b) comparison of brain metabolism of non-toxic model, and toxic model under 100 times steady state toxin fluxes, (c) comparison of sIEC metabolism of non-toxic model, and toxic model under steady state toxin fluxes, and (d) comparison of SIEC metabolism of non-toxic model, and toxic model under 100 times steady state toxin fluxes

Table S5: Table describing the applications of pareto optimization. (a) Unweighted pareto optimization cases, (b) Weighted pareto optimization cases.

Table S6: Table describing the comparison of models under different constraints, and the number of reactions affected. Default diet is western diet if not mentioned otherwise.

Table S7: Excel file containing details of the individual bacterial products. Minimum and maximum fluxes (mmol/gDW/hr) of the secretion reactions of: *Bacteroides Vulgatus, Desulfovibrio desulfuricans, Clostridium perfringens, Lactobacillus acidophilus and Bifidobacterium longum*

Table S8: Excel file containing details of the effect of adding beneficial bacteria percentages simulated on the integrated model, as toxin steady state concentrations (M) as a function of beneficial bacteria percentage, under high-fibre diet, toxin steady state fluxes (mmol/gDW/hr) as a function of beneficial bacteria percentage, under high-fibre diet, toxin steady state concentrations (M) as a function of beneficial bacteria percentage, under western diet, and toxin steady state fluxes (mmol/gDW/hr) as a function of beneficial bacteria percentage, under western diet

Table S9: Excel file containing details of effect of bacteria and diet on gut metabolism, i.e., effect of probiotics on sIEC metabolism, effect of harmful bacteria on sIEC metabolism, and effect of changing diet from western to high-fibre diet on SIEC metabolism

Table S10: Table containing details of superoxide SOX Steady state Concentrations (M)

Table S11: Excel file containing details of the integrated model results, i.e., toxin steady state concentrations (M) and toxin steady state fluxes (mmol/gDW/hr)

Table S12: Excel file containing details of the bacterial secretion products, i.e., harmful bacteria secretions ranked from highest to lowest based on maximum fluxes, reported in mmol/gDW/hr, beneficial bacteria secretions ranked from highest to lowest based on maximum fluxes, reported in mmol/gDW/hr, effect of changing diet from western to high-fibre diet, and effect of addition of probiotics to a gut microbiome with harmful bacteria secretion products

Table S13: Excel file containing details of the diet constraints and intestinal conditions.

Table S14: Table detailing the gut - microbiome exchanges Table S15: Table for steps involved in PBPK modelling

Table S16: Table for the symbols and subscripts used in the mathematical representation of the PBPK model

Table S17: Table containing the characteristic of each compartment in PBPK model

Table S18: Table containing the details of the physiological parameters used in the PBPK model

Table S19: Table containing the values of kinetic parameters used in the metabolism terms for the PBPK model

Table S20: Table containing the train set matrix size for each tissue type used in the PBPK model

